# Perfusion Bioreactor Culture Incorporating Mechanical Confinement Enhances Mesenchymal Stem Cell Extracellular Vesicle Production and Wound Healing Potential

**DOI:** 10.1101/2025.08.12.669872

**Authors:** Emily H. Powsner, Stephanie M. Kronstadt, Nicholas H. Pirolli, Talia J. Solomon, Raith Nowak, Kristin Nikolov, William Pieper Holeman, Ian M. Smith, Kimberly M. Stroka, John P. Fisher, Steven M. Jay

## Abstract

Mesenchymal stem cell extracellular vesicles (MSC EVs) have been widely studied for regenerative medicine and tissue repair applications. However, clinical translation of EV therapeutics has been hampered by low potency and lack of scalable production strategies. This work aims to develop a novel approach that exploits the mechanosensitivity of MSCs to enhance EV potency in the context of enhanced production via bioreactor culture. MSCs are well known to respond to mechanical stimuli such as substrate stiffness and shear stress, and here it is shown that exposing MSCs to another mechanoregulatory parameter, confinement, enhances the pro-angiogenic bioactivity of their EVs. This is consistent across both donor-derived primary MSCs and induced pluripotent stem cell-derived MSCs (iMSCs). The development of a 3D-printed perfusion bioreactor system that enables culture of confined MSCs under flow is also detailed here, resulting in a 67-fold increase in EV production compared to flask culture. iMSC EVs obtained downstream of this confinement-bioreactor culture induce greater vascularization and generally improve wound healing in a diabetic mouse model compared to iMSC EVs from conventional tissue culture. Overall, this work establishes the development of a scalable, bioreactor-based iMSC EV production platform that provides a solution to major translational bottlenecks of therapeutic EVs.

## INTRODUCTION

Extracellular vesicles (EVs) have been established as critical mediators of cell-to-cell communication [1]. Their ability to carry and transfer cargo that mirrors that of their producer cells makes stem cell-derived EVs an attractive cell-free alternative to conventional stem cell therapies [2]. In particular, mesenchymal stem cell (MSC) EVs have been recognized for their ability to promote vascularization and tissue repair and modulate the immune system, while mitigating cell-associated risks of tumorigenicity, spontaneous differentiation, and immunogenicity [2–4]. However, FDA approval and commercialization of EV therapies have not correlated with the exponential growth of pre-clinical EV research or clinical trials [5].

In terms of the clinical translation of EVs, low innate potency (which necessitates higher and/or multiple doses), donor variability (which contributes to scalability and quality control issues), and a lack of scalable manufacturing methods all remain major obstacles [6, 7]. Donor bone marrow-derived MSCs (BMMSCs), which are the most common source for MSC EVs [5], are substantially limited by passage number and donor differences in efficacy [8, 9]. While MSC immortalization holds the potential to overcome passage limitations and donor variability, the introduction and permanence of oncogenes in this process poses poorly understood risks of genetic mutation and the potential for the EVs to induce tumorigenesis in recipient cells [10]. As a solution to low innate potency, genetic engineering and exogenous therapeutic cargo loading have been employed and have demonstrated efficacy, especially at the pre-clinical scale [11–13]. However, both approaches are often time consuming and costly processes that sacrifice EV integrity, require intensive sample purification [14].

It is well established that cells can sense and respond to mechanical stimuli from their microenvironment, such as substrate stiffness and shear stress [15–20]. Another mechanoregulatory phenomenon of interest that has been less studied, specifically with regards to EV production is cellular confinement, particularly that which forces an elongated morphology. One study showed a relationship between morphological elongation of MSCs in response to IFN-*γ* stimulation and their immunosuppressive capacity [21], and there is evidence that confinement impacts MSCs’ secretome [15]. Consequently, we expect that confining MSCs could affect the production and therapeutic capacity of their EVs in a scalable manner.

Accordingly, we studied the influence of cellular confinement on the reparative, pro-angiogenic effects of MSC EVs for wound repair. To assess potential therapeutic impacts of these phenomena, we innovated upon a previously published perfusion bioreactor our group developed that enabled an 83-fold increase in BMMSC EV production [8]. Induced pluripotent stem cell-derived MSCs (iMSCs) were used as EV producer cells given their ability to overcome donor variability and senescence challenges. Our results show that mechanical confinement results in increased iMSC EV bioactivity relevant to wound repair. Further, this effect can be maintained in perfusion bioreactor culture in tandem with an increase in EV production. Overall, the results address several of the major obstacles to clinical translation of therapeutic MSC EVs for wound repair.

## RESULTS

### Mechanical confinement of MSCs augments EV bioactivity

For initial *in vitro* studies, the mechanical confinement of cells was induced using PDMS-based micropillar arrays with pillars 5 or 50 µm apart based on devices developed by Doolin et. al. [22] (Figure 1A, B). PDMS with no pillars and collagen-coated flasks were used as controls, and the expected morphological changes were confirmed visually (Figure 1C). Isolated iMSC EVs were confirmed to have the expected size (Figure S1A), characteristic EV markers (Figure S1B), and morphology (Figure S1C). Angiogenic activity was evaluated with a tube formation assay using human umbilical vein endothelial cells (HUVECs), where iMSC EVs from 5 µm-confined producer cells exerted a significantly greater angiogenic effect compared to EVs from 50 µm-confined cells, and even more so compared to EVs from cells cultured under the no pillar condition as well as flask-cultured cells (Figure 2A). A similar trend was observed when measuring migration of iMSC EV-treated HUVECs, where the EVs’ ability to induce cell migration was proportional to the degree of confinement (Figure 2B). Anti-inflammatory activity of the EVs was also conserved, but not further improved, as IL-6 and TNF-⍰ expression by RAW 264.7 mouse macrophages treated with all EV groups was suppressed (Figure S2).

**Figure 1:**
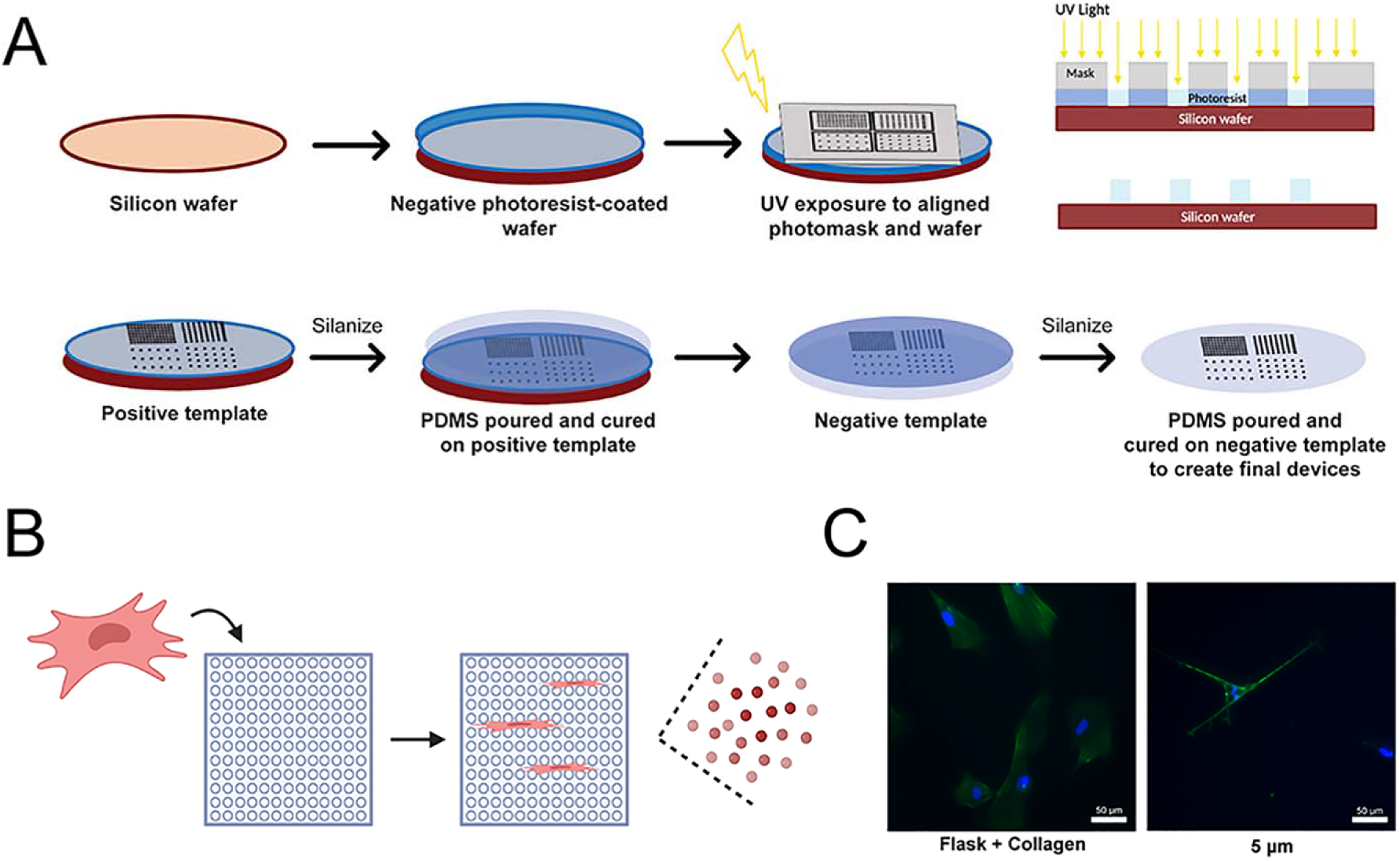
Confinement device fabrication. A) Schematic to illustrate the steps to fabricate 36 mm x 36 mm micropillar devices with photolithography and PDMS. B) Simplified workflow to isolate EVs from confined cells. iMSCs are seeded on micropillar arrays and left to adhere overnight before changing to EV-depleted media and subsequently collecting the conditioned media the following day. C) Representative merged fluorescence microscopy images taken at 24x magnification of Hoechst- and phalloidin-stained iMSCs in flask culture and 5 µm micropillar device culture. Green shows phalloidin staining and blue shows Hoeschst staining. Scale bars = 50 µm.

**Figure 2:**
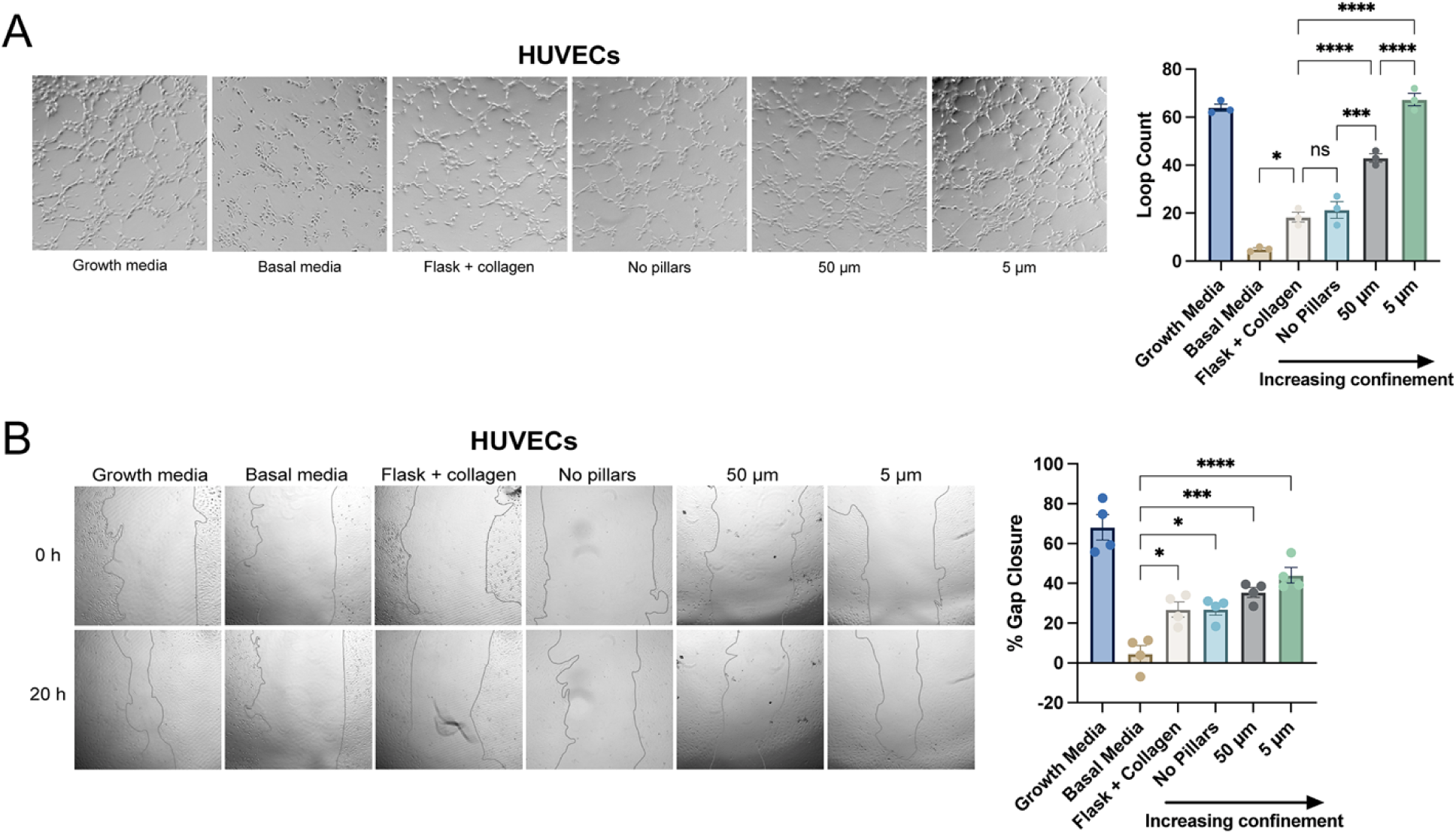
iMSC EV bioactivity is responsive to producer cell confinement. A) Tube formation assay data quantified by loop count of HUVECs treated with either endothelial growth media, endothelial basal media, or 5×10^9^ EVs mL^-1^ from flask culture, no pillar PDMS culture, 50 µm-confined culture, or 5 µm-confined culture, and the corresponding representative images (n=3). B) Percent gap closure of HUVECs treated with endothelial growth media, endothelial basal media, or 5×10^9^ EVs mL^-1^ from flask culture, no pillar PDMS culture, 50 µm-confined culture, or 5 µm-confined culture, and the corresponding representative images (n=3). All data is representative of 3 independent experiments. All values expressed as mean +/- standard error of the mean (ns – no significance; * p<0.05; *** p<0.001; **** p<0.0001).

These angiogenic and anti-inflammatory effects of confinement were validated by also evaluating EVs from BMMSCs, which is a more established cell source compared to iMSCs, albeit with limitations related to donor variability and senescence. BMMSCs were seeded within 5 and 50 µm confinement devices, as well as in flasks and no pillar PDMS devices in the same manner as the iMSCs. Isolated EVs were used to treat HUVECs in both gap closure and tube formation assays, where EVs from cells exposed to greater degrees of confinement induced a more migratory and pro-angiogenic effect compared to flask EVs and basal media (Figure S3), reflecting the trends observed with iMSC EVs.

### Incorporation of MSC confinement within a perfusion bioreactor enhances EV production with retained bioactivity

While inducing confinement improved the angiogenic potential of iMSC EVs, EV yield remained relatively low and comparable with flask culture, with a non-significant trend of increased EV production per cell. It has been well established that bioreactors are a scalable method to effectively culture cells and in particular, our group has reported that a perfusion bioreactor can increase bone marrow-derived MSC EV yield by 83-fold compared to flask culture [8]. Thus, we redesigned the previous 3D-printed perfusion bioreactor to allow for the incorporation of 5 µm PDMS micropillar arrays given the evident impact on bioactivity (Figure 3A).

**Figure 3:**
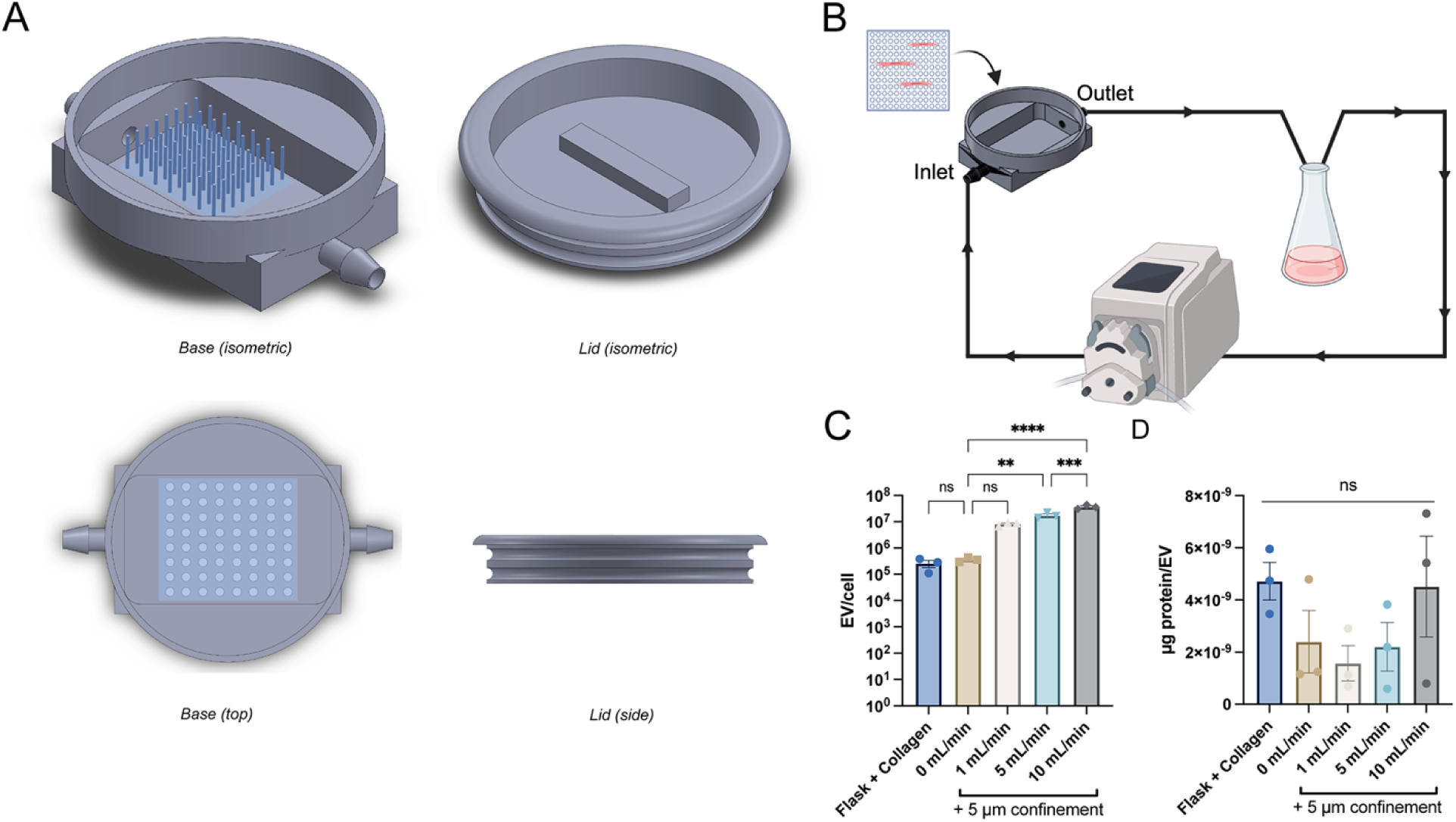
Perfusion bioreactor culture incorporating cell confinement enhances iMSC EV yield. A) SolidWorks renderings of the 3D-printed perfusion bioreactor, with the inserted PDMS micropillar array, and lid, which is sealed using an O-ring. The internal chamber where media is able to fill is 64 mm x 40 mm x 10 mm. The pillared array is not represented at scale to better visualize the pillars. Actual pillars are 5 µm apart and ∼14-17 µm high. B) Schematic of the bioreactor-pump setup. The seeded PDMS micropillar device is inserted into the bioreactor, the lid is placed, and then bioreactor is connected to a peristaltic pump, with the ends of the tubing placed into a 70 mL media reservoir. B) iMSC EV production quantified as EV per cell exposed to bioreactor flow rates of 1, 5, and 10 mL min^-1^ with static confinement culture and flask culture as controls. Data shows the average of 3 independent experiments (n=3). C) Protein per EV from 1, 5, and 10 mL min^-1^ bioreactor flow rates with static confinement and flask culture as controls. Data shows the average of 3 independent experiments (n=3). All values expressed as mean +/- standard error of the mean (ns – no significance; ** p<0.01; *** p<0.001; **** p<0.0001).

We then sought to optimize this confinement-bioreactor system for both maximum cell retention and EV yield using the illustrated setup (Figure 3B). Testing flow rates of 1, 5, and 10 mL min^-1^, we found that iMSC EV production per cell could be increased by 32-, 67-, and 140-fold, respectively compared to flask culture (Figure 3C). We also measured protein per EV as a proxy for sample purity, where exposure to 1 and 5 mL min^-1^ flow rates resulted in a 3- and 2-fold decrease in protein content, respectively, while the 10 mL min^-1^ flow rate produced samples with the same average protein concentration as flask culture (Figure 3D). Based on these data, we determined that 5 mL min^-1^ was optimal for increasing EV yield, ensuring cell retention within the device, and improving purity.

Importantly, we next confirmed that the amplified angiogenic bioactivity of the EVs that was observed with cell confinement was not compromised by flow (Figure 4A). Given that it was not, we continued to move forward with a 5 mL/minute flow rate. We directly compared the bioactivity of EVs from iMSCs grown on collagen-coated flasks (hereinafter referred to as ‘flask’), EVs from 5 µm-confined iMSCs without flow (‘static’), and EVs from 5 µm-confined iMSCs exposed to 5 mL min^-1^ flow in the perfusion bioreactor (‘bioreactor’). With the goal of assessing bioactivity in cell types directly associated with wound healing, we started with human dermal microvascular endothelial cells (HDMECs). We found that treatment with both static- and bioreactor culture-derived EVs significantly increased tube formation compared to flask EVs (Figure 4B). The same trend was observed with fibroblast migration, where the static- and bioreactor culture-derived EVs exerted a statistically significantly higher gap closure effect than the flask culture-derived EVs (Figure 4C). Although the static and bioreactor EVs caused slightly more keratinocyte migration, it was not statistically significant compared to the flask EVs (Figure 4D). All of the EVs were able to reduce pro-inflammatory cytokine secretion in RAW 264.7 mouse macrophages at similar levels, with the bioreactor culture-derived EVs showing a slightly more potent effect in suppressing IL-6 expression (Figure S4).

**Figure 4:**
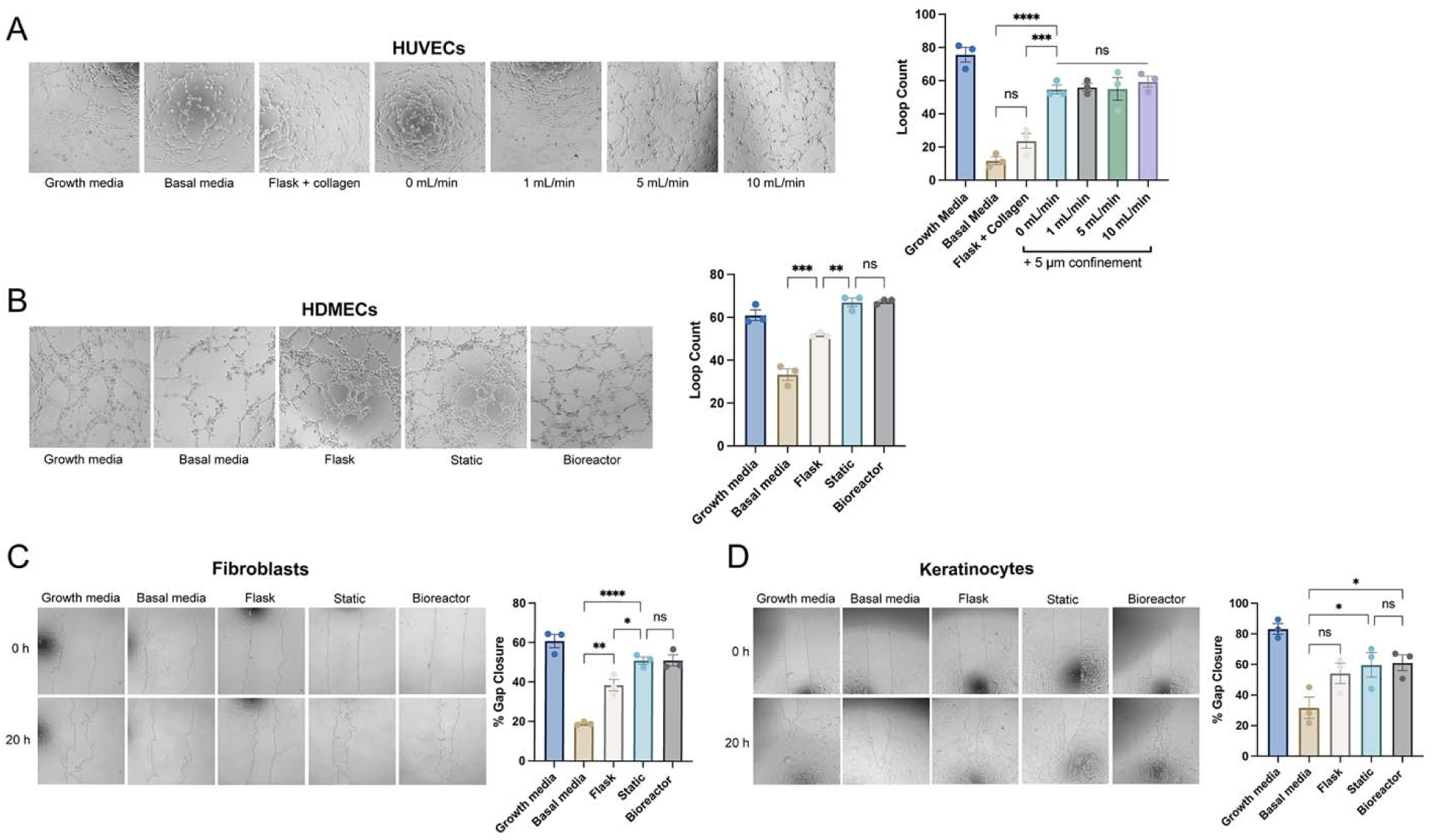
Improved iMSC EV wound healing activity is maintained in the confinement-bioreactor system. A) Tube formation by HUVECs treated with endothelial growth and basal media or 5×10^9^ EVs mL^-1^ from flask culture or bioreactor culture with 1, 5, and 10 mL min^-1^ flow rates, and the corresponding representative tube formation images (n=3). B) Tube formation by HDMECs treated with endothelial growth or basal media or 5×10^9^ EVs mL^-1^ from flask culture, static 5 µm confinement culture, or 5 mL min^-1^ + 5 µm confinement bioreactor culture, and representative images (n=3). C) Percent gap closure of human dermal fibroblasts after 20 hours with treatment of fibroblast growth or basal media or 5×10^9^ EVs mL^-1^ from flask culture, static 5 µm confinement culture, or 5 mL min^-1^ + 5 µm confinement bioreactor culture, and representative images (n=3). D) Percent gap closure of human epidermal keratinocytes after 20 hours with treatment of keratinocyte growth or basal media or 5×10^9^ EVs mL^-1^ from flask culture, static 5 µm confinement culture, or 5 mL min^-1^ + 5 µm confinement bioreactor culture, and representative images (n=3). All data is representative of 3 independent experiments. All values expressed as mean +/- standard error of the mean (ns – no significance; * p<0.05; ** p<0.01; *** p<0.001; **** p<0.0001).

### EVs from confined iMSCs in perfusion bioreactor culture improve wound repair in a diabetic mouse model

To determine if the results we observed *in vitro* translated *in vivo* in a more complex model, we utilized a splinted diabetic mouse wound model (Figure 5A). Injections were administered on days 0, 3, 6, and 9, and wounds were imaged to measure wound closure throughout the healing process. The only statistically significant difference in percent closure was observed on day 14 between the PBS-treated and bioreactor EV-treated mice (Figure 5B, C). Interestingly, wound closure improved for the flask and bioreactor EV-treated groups, while it initially worsened for the PBS-treated mice on day 3 (Figure 5C). A faster rate of wound closure with less variation between mice was also observed for the bioreactor EV-treated group, especially between days 0 and 9. From the wound tissue harvested on day 16, qPCR for M1 macrophage markers indicated a slight decrease in the bioreactor EV-treated group compared to the flask EV-treated group (Figure 5D). However, only iNOS achieved an average expression level below that of the PBS-treated group with treatment with bioreactor EVs. M2 macrophage markers were at roughly the same level between flask and bioreactor EV treated groups, with at least a 2-fold increase in IL-10 and Arg1 in the bioreactor EV group compared to the PBS group (Figure 5E). Upon analyzing the tissue architecture, the amount of regenerating epithelium (as a percentage of the total wound bed length) (Figure 5F), collagen deposition via Masson’s Trichrome staining (Figure 5G), and granulation tissue area (Figure 5H) revealed no statistically significant differences between groups, although the granulation tissue area trended higher in the EV-treated groups compared to the PBS-treated mice. Immunohistochemistry staining for CD31, a marker of blood vessel formation, while not statistically significant, showed a 2.4- and 1.4-fold increase in CD31 intensity normalized to DAPI in the bioreactor group compared to the flask and PBS groups, respectively (Figure 5I).

**Figure 5:**
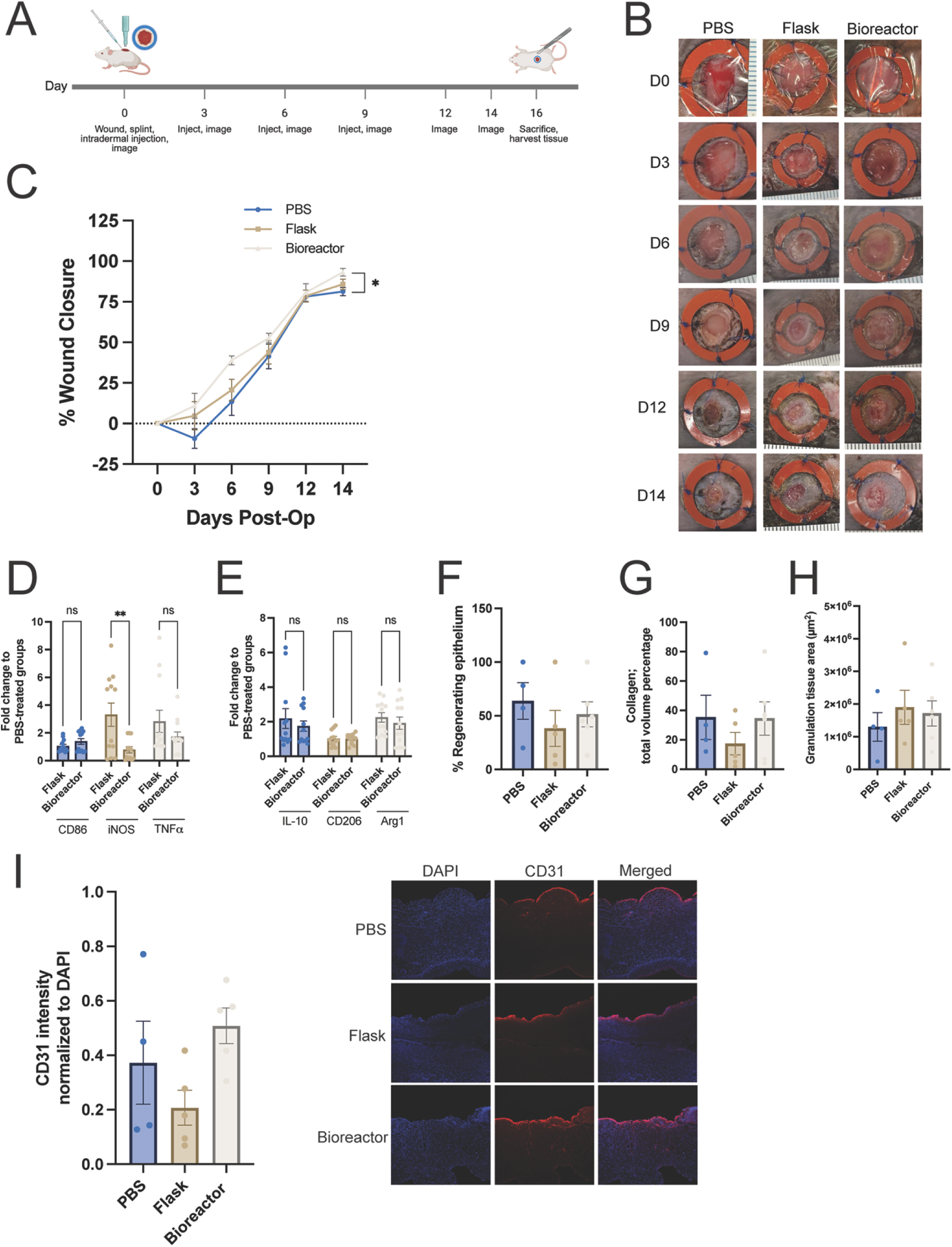
Bioreactor EVs improve tissue repair in a diabetic mouse wound model. A) Timeline of experiment setup and injection schedule. Mice received injections of 1.1×10^10^ EVs or an equivalent volume of PBS on days 0, 3, 6, and 9. B) Representative photos of splinted wounds receiving each treatment over 14 days (n=4-6). C) Percent wound closure based on wound area over 14 days (n=4-6). D) Fold change of M1 markers CD86, iNOS, and TNF11 and E) M2 markers IL-10, CD206, and Arg1 in harvested tissue from mice receiving flask and bioreactor EV injections relative to tissue from mice receiving PBS, as quantified by qPCR. Data represents 4 biological replicates analyzed with 3 technical replicates each per treatment group (n=4). F) Length of regenerating epithelium from H&E staining as a percentage of the length of the wound bed (n=4-6). G) Collagen deposition from Masson’s Trichrome staining quantified as a percentage of the total tissue volume (n=4-6). G) Granulation tissue area from H&E staining (n=4-6). I) CD31 fluorescence intensity normalized to DAPI over multiple fields of view and representative confocal microscopy images taken at 10X magnification (n=4-5). All values expressed as mean +/- standard error of the mean (ns – no significance; * p<0.05; ** p<0.01).

### Mechanical confinement affects the molecular cargo of iMSC EVs

From a therapeutic standpoint, proteins and RNAs are key components of EVs. miRNAs play multifaceted roles in both physiological and pathological processes, and a single miRNA can target multiple signaling pathways, making them potent effector molecules [23]. They are also frequently implicated as mediators of the therapeutic effects conferred by EVs [24–27]. Proteins are also critical components of EVs in the context of both biogenesis and as transcription factors and signaling molecules that can target intracellular signaling pathways [28–30]. Thus, to examine the effects of confinement and flow on EV cargo, we compared the miRNA profiles between EVs from each of the different culture conditions (i.e. flask, static, bioreactor) using an MSC EV miRNA qPCR array. A total of 166 miRNAs associated with MSC EVs were probed for, with 68 detected between at least two of the biological replicates (Figure 6A). Compared to the flask EVs, 15 were downregulated and 34 were upregulated more than 1.5-fold in the static EVs, and 20 were downregulated and 36 were upregulated more than 1.5-fold in the bioreactor EVs (Figure 6B, C). Notably, there were only three miRNAs that were statistically significantly differentially expressed in both the static and bioreactor EVs: let-7e-5p, miR-182-3p, and miR-221-3p.

**Figure 6:**
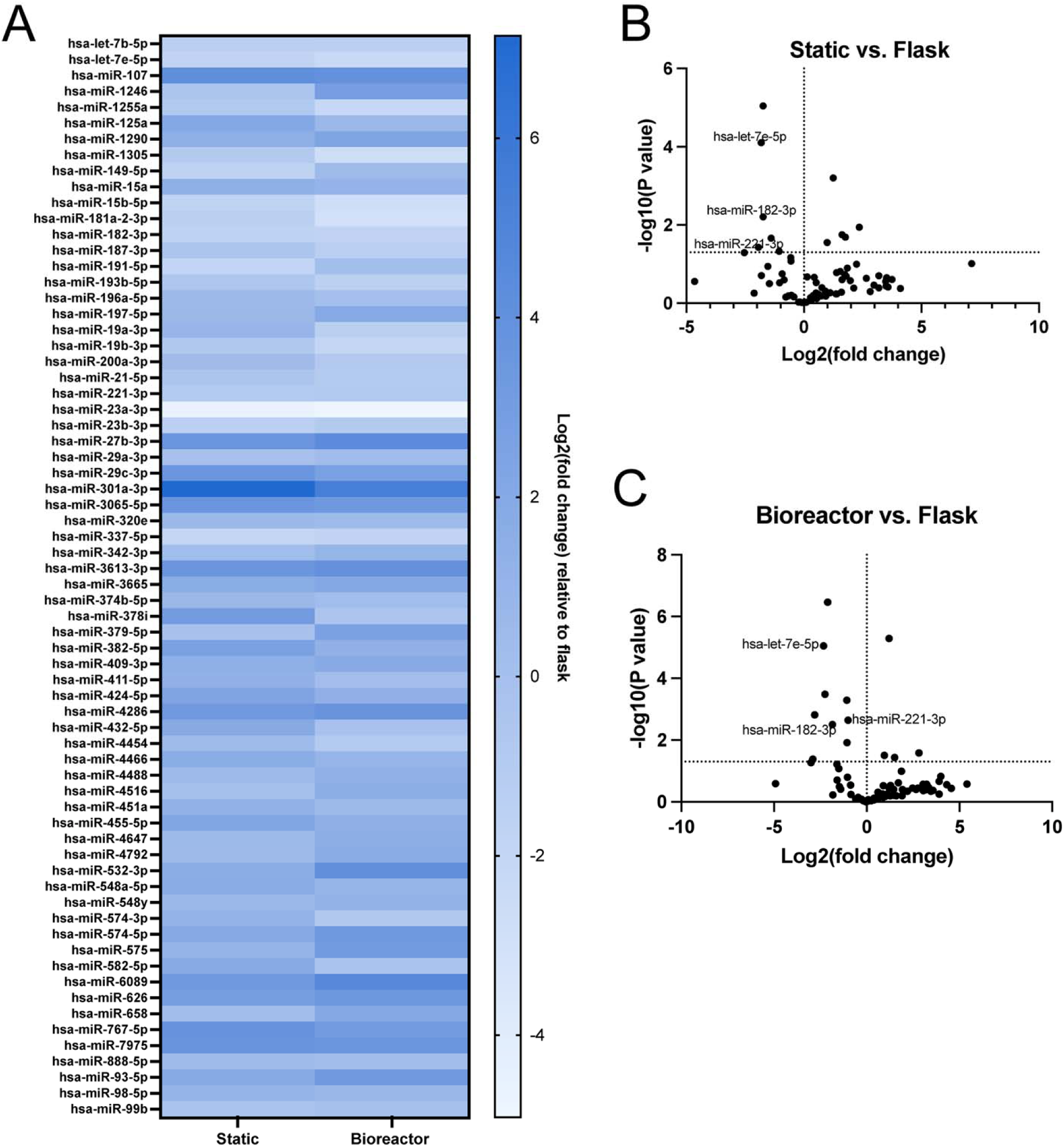
Comparison of miRNA profiles between iMSC EVs from different culture conditions. A) Heat map of the miRNA content of 5 µm static and 5 µm bioreactor EVs represented as the log2(fold change) relative to flask EVs (n≥2). B) Volcano plot comparing the miRNA within static and flask EVs, with the horizontal line at 1.3 (p = 0.05) representing statistical significance (n≥2). C) Volcano plot comparing the miRNA within bioreactor and flask EVs, with the horizontal line at 1.3 (p = 0.05) representing statistical significance. The labeled points indicate those in common between both static and bioreactor EVs (n≥2). All data represents the average of at least 2 independent experiments.

We also compared the protein content of all three EV groups to identify differences that may be influenced by mechanical stimuli and the cell culture conditions. Across all groups, there were 52 proteins in common with a significance score over 20 (p > 0.01) based on label-free quantification methods in the PEAKS software (Figure 7A). When comparing protein abundance between bioreactor and flask EVs, 27 were downregulated in the bioreactor group while 25 were upregulated (Figure 7B). Interestingly, when comparing to the static EVs, all proteins except for 3 were upregulated (Figure 7C). Gene ontology and pathway enrichment analysis of all upregulated proteins in the bioreactor EVs relative to flask EVs was also performed [31–33]. Of the 20 biological processes with the lowest p-values (p ≤ 0.02), which indicated the highest probability of the protein being associated with the specific pathway/process, many are directly or indirectly involved in wound healing (Figure 7D).

**Figure 7:**
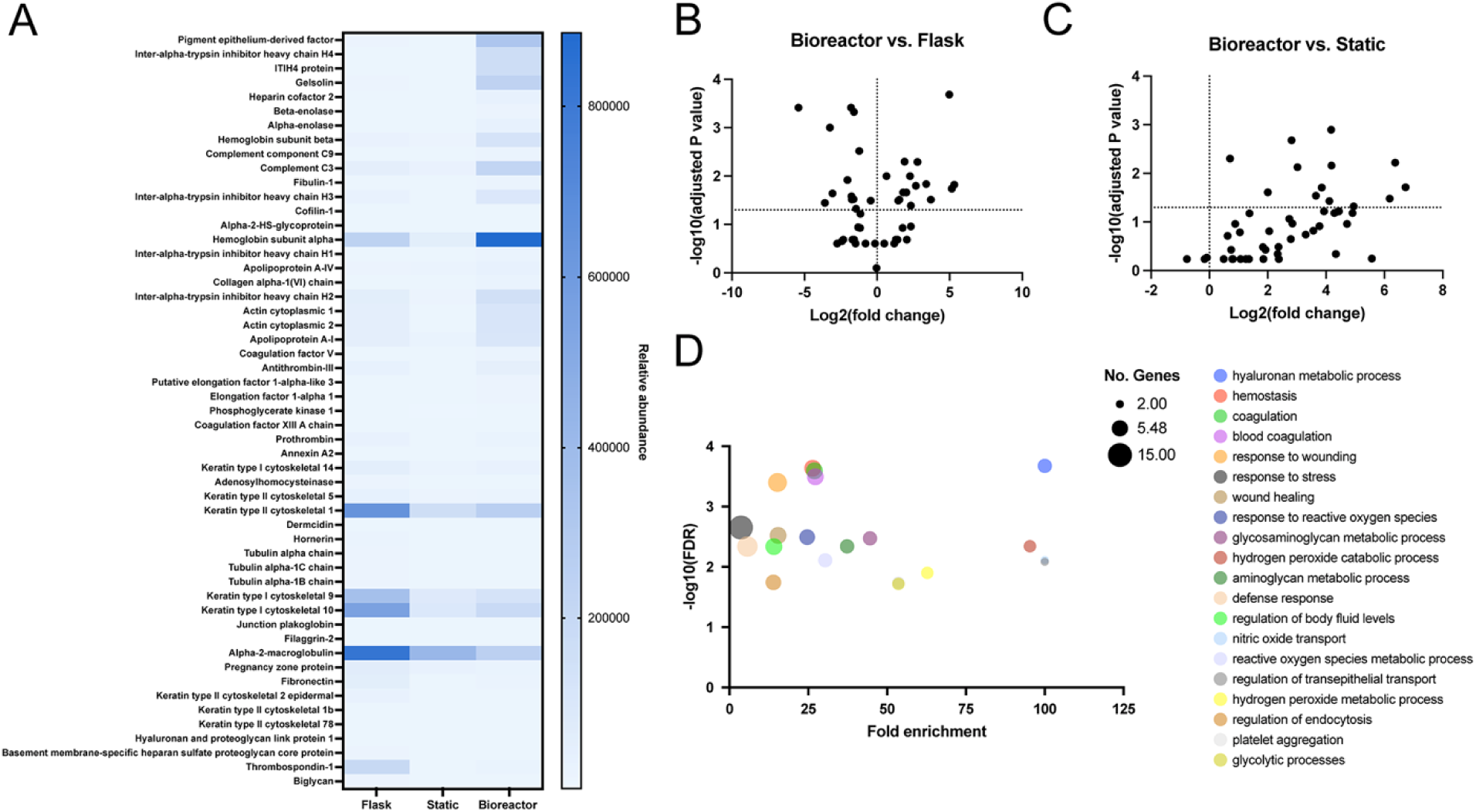
Proteomics analysis for iMSC EVs from different culture conditions. A) Heat map of the relative abundance of protein content of flask, static, and bioreactor EVs from proteomics analysis (n=2). B) Volcano plots comparing the protein in bioreactor and flask EVs, with the horizontal line at 1.3 indicating statistical significance (p=0.05) (n=2). C) Volcano plots comparing the protein in bioreactor and static EVs, with the dotted line at 1.3 indicating statistical significance (p=0.05) (n=2). D) Gene ontology biological process enrichment analysis of all upregulated proteins in the bioreactor EVs compared to flask EVs. Fisher’s exact test with false discovery rate correction (FDR) was used to generate the adjusted p-value, and the processes with the 20 smallest p-values were plotted. Data represents the average of 2 technical replicates.

## DISCUSSION

Clinical translation of EV therapeutics, despite demonstrated pre-clinical efficacy, has been severely hindered by a myriad of issues related to potency, scalable manufacturing platforms, and cell source and donor variability [6, 34]. Mechanical cues, specifically the physical confinement of cells, can induce phenotypic changes [21, 35]. As such, we investigated how confinement and flow affects the production and therapeutic efficacy of MSC EVs through the development of a 3D-printed perfusion bioreactor system incorporating cell confinement. To overcome donor cell-specific challenges faced with BMMSCs, iMSCs were used in these studies to bolster their plausibility as a renewable MSC EV source with reduced donor variability and to establish their compatibility with this scalable EV manufacturing platform.

Perhaps the most widely studied approach to improving potency is the overexpression of RNAs in the producer cells for subsequent trafficking into EVs [11, 36]. From a translational standpoint however, genetic engineering of cells requires substantially more time, money, and optimization upstream and more intensive purification downstream. Doolin et al. demonstrated that confinement of MSCs can effectively influence cell behavior [15, 37]. Therefore, we sought to identify if this mechanical cue of confinement, which forces the cells into an elongated spindle shape, could increase iMSC EV potency and found that the effects on phenotype and angiogenic function indeed translated to their EVs (Figure 2). Huang et al. also described a correlation between cell elongation via alignment, and bioactivity - but with macrophages - where polarization to an M2 phenotype was induced and the angiogenic and osteogenic capacity of the EVs was augmented upon forcing this confined shape [38]. Overall, this approach allowed us to generate more potent EVs while changing only the culture substrate - the PDMS micropillar array - without the use of extraneous reagents, chemicals, and/or cargo.

Despite our results showing confinement-induced increases in potency, confined cells did not produce a significantly greater amount of EVs compared to flask-grown cells. Hence, we redesigned a 3D-printed perfusion bioreactor that was previously shown by our group to be biocompatible and effective at increasing BMMSC EV production [8] to now incorporate cell confinement (Figure 3A). Based on quantification of protein per EV, which is commonly used as a proxy for purity [39], and EV production per cell, we concluded that a 5 mL min^-1^ flow rate was optimal, with a significant 67-fold increase in EV production and a 2-fold increase in purity compared to flask culture (Figure 3C, D). These findings were consistent with other reports where fluid flow was used as an effective strategy to increase EV yield, although such systems did not incorporate additional mechanical cues. For instance, Costa et al.’s stirred tank bioreactor setup with MSCs resulted in ∼3-fold increase in EVs per cell compared to static flask culture [40]. Patel et al. achieved a ∼100-fold increase in total HDMEC EV yield with perfusion bioreactor culture compared to static culture, although these data did not appear to be normalized to cell count at the time of collection [41]. Our group also reported an 83-fold increase in MSC EVs per cell compared to flask culture using a perfusion bioreactor [8] that served as the precursor to the system used in these studies. Accordingly, flow-derived shear stress is known to increase membrane tension, blebbing, and exocytic activity, which are modes of EV biogenesis [42, 43]. However, while all of these bioreactor systems led to significant increases in EV production, there were no significant differences in *in vitro* bioactivity reported between static culture- and bioreactor-generated EVs across the respective studies.

Therefore, we examined the *in vitro* angiogenic and wound repair bioactivity of the EVs produced using our system, looking at HDMECs, fibroblasts, keratinocytes, and macrophages; all cells that play critical roles in the wound healing process, and also exhibit dysfunction either because of or leading to chronic wounds [44–49]. Consequently, the ability to promote angiogenesis and cell migration in all three skin cell types is imperative for an effective therapeutic for chronic wound healing. The ability for a therapeutic to reduce inflammation would also prove beneficial in helping overcome the prolonged inflammatory state seen in chronic wounds, which all EV groups were able to achieve *in vitro* (Figure S2, S4). While the bioreactor EVs showed no difference in angiogenic bioactivity compared to the static EVs across all skin cell types and both assays, both groups exerted a significantly greater effect compared to the no treatment (basal media) control and to the flask EVs, allowing for significantly fewer EVs per dose to achieve the same effect (Figure 4). This was a critical finding, as we have established that not only can we improve yield, but also potency of the EVs simultaneously using our bioreactor system.

Given that wound healing is a complex and dynamic process, we next evaluated the efficacy of these EVs *in vivo* to assess their effects on wound healing more holistically. The diabetic (db/db) mouse wound model is widely used to test treatments for wound healing, including various EV formulations developed by our group and others [8, 11, 50, 51]. However, one major drawback of this model is that mouse skin heals primarily by contraction while human skin heals primarily by re-epithelialization, making the treatment effects often difficult to discern from their natural healing process [52]. Therefore, we utilized a silicone splint to minimize contraction and promote a granulation and re-epithelialization process that more closely mimics that of humans [52]. All treatments, including the vehicle control of PBS, achieved a similar level of wound closure by day 14 with statistical significance between the PBS and bioreactor EV treatment groups (Figure 5C). Interestingly, the biggest average differences in wound closure occurred on days 3 and 6, where the PBS-treated wounds increased in area on day 3. This observation is consistent with other groups also using db/db mouse wound healing models [11, 53], and aligns with the physiological process of wound healing where peak inflammation occurs about 3 days post-wounding [54]. However, the EV treatments, especially the bioreactor EVs, were able to negate this initial healing delay and promote a generally faster rate of wound closure than PBS or flask EVs between days 0 and 9. Outside of overall closure, tissue-level and molecular-level analysis revealed few differences between groups (Figure 5D-H). It is important to note that the tissue was harvested and analyzed from mice at day 16. This endpoint was chosen mainly to analyze vascularization and healed tissue architecture since the differences between treatments *in vitro* were pronounced only in the angiogenesis assays and not in the anti-inflammatory assays. However, since peak inflammation occurs at an earlier time point, inflammatory marker expression and treatment effects on inflammation would likely be more accurately and reliably measured around days 3-6, as was done by Levy et al. [50]. We did observe a slight increase, albeit not statistically significant, in vascularization with the bioreactor EV treatments by the increase in CD31+ blood vessel formation observed in wound tissue from the bioreactor EV-treated mice (Figure 5I). While results using similar models seem to vary between studies, one group examined EV-treated wound tissue histologically on day 7 and 14, where both CD31 and collagen deposition showed more dramatic differences between the control and EV-treated groups on day 7, further supporting it as a time point worth investigating more closely in the future [55]. We also stopped treatment after day 9, which is when the differences in the rate of closure decreased, so continuing to treat through the entirety of the study (i.e. day 12, 14) may also help discern other differences between treatments. Other groups such as Yu et al. used a slightly different approach where they mixed MSC EVs with an injectable hydrogel to treat diabetic wounds, reporting that the combination improved blood vessel formation and overall wound healing more so than the EVs alone [56]. Similarly, Xiao et al. showed that compared to MSC EVs, a decellularized scaffold loaded with EVs was able to significantly increase collagen deposition and CD31 blood vessel formation [55]. These results suggest that our EVs’ efficacy may be able to be further improved by containing them at the wound bed using a biocompatible scaffold, which could be an interesting future avenue.

As with any study, specifically with the inherent variability that comes with animal models, there are limitations. There is no standardized dosing scheme established for EVs or for this model, which not only impacts therapeutic efficacy but also makes it difficult to compare results between studies. Here, we maximized the dose based on EV yields to help ensure consistency and reduce artifacts from batch-to-batch variation. Notwithstanding practical limitations regarding dosing schedule, batch yields, and cohort size, a better understanding of EV dosing through dose escalation studies would be helpful to optimize this model in the future.

Nevertheless, it is clear that both static confinement and confinement in bioreactor culture impacts the production of iMSC EVs functionally, and at a molecular cargo level, as we observed the up and downregulation of various miRNAs and proteins within the vesicles (Figure 6, 7). The upregulated proteins in the bioreactor EVs when compared to the flask EVs also had significant enrichment in many biological processes including wound healing and those related to wound healing including hemostasis, coagulation, and nitric oxide transport (Figure 7) [44, 57]. The numerous changes in cargo expression also suggest that the uptick in bioactivity is likely attributed to multiple molecules and mechanisms working synergistically. While these analyses are limited and presented merely as observations that would require mechanistic studies to understand their implications, they provide valuable information that could be used as a starting point for future studies to delve deeper into specific mechanisms of action or to identify potential critical quality attributes. A bioinformatics approach would also be helpful to more thoroughly analyze such data in the future.

Overall, we have developed and optimized a novel EV production platform utilizing mechanobiology and a 3D-printed perfusion bioreactor to simultaneously increase the yield and potency of iMSC EVs for wound repair. Specifically, we showed that bioactivity was enhanced with the mechanical cue of cell confinement and retained when moved into bioreactor culture, which was able to increase production 67-fold. In doing so, we were able to overcome key bottlenecks to clinical translation and improve the scalability of iMSC EVs with this relatively simple and cost-effective approach that could likely be adapted for other mechanoresponsive cell and EV types. Despite the array of additional studies that could be performed to better understand MSC EVs for wound healing specifically, the results here provide a basis for and strongly support the potential of this manufacturing platform for scaling up potent iMSC EV production.

## METHODS

### Cell culture

Human induced pluripotent stem cell-derived MSCs (iMSCs) and human bone marrow-derived MSCs (BMMSCs) were purchased from ATCC (ACS-7010, PCS-500-012) and cultured in Dulbecco’s Modified Eagle’s Medium (DMEM) (Corning; 10-013-CV) supplemented with 10% fetal bovine serum (FBS) (Cytiva; SH30910.03), 1% penicillin-streptomycin (VWR; 45000-652), and 1% MEM non-essential amino acids (ThermoFisher Scientific; 11140050).

Human umbilical vein endothelial cells (HUVECs) and human dermal microvascular endothelial cells (HDMECs) were purchased from PromoCell (C-12203, C-12212) and cultured in endothelial growth media (PromoCell; C-2212) with 1% penicillin-streptomycin in 0.1% gelatin-coated tissue culture flasks.

Human primary epidermal keratinocytes were purchased from ATCC (PCS-200-011) and cultured in dermal cell basal media (ATCC; PCS-200-030) supplemented with ATCC’s Keratinocyte Growth Kit components (ATCC; PCS-200-040) and 0.1% penicillin-streptomycin.

Human dermal fibroblasts were purchased from ATCC (PCS-201-012) and cultured in Gibco DMEM/F-12 (ThermoFisher Scientific; 11320033 supplemented with 10% FBS and 1% penicillin-streptomycin.

RAW 264.7 mouse macrophages were purchased from ATCC (TIB71) and cultured in DMEM supplemented with 5 % FBS and 1% penicillin-streptomycin.

### Micropillar device fabrication, preparation, and seeding

Micropillar devices were made using polydimethylsiloxane (PDMS) (Krayden; DC2065622) and photolithography as described previously [22]. Briefly, a photo mask was made using AutoCAD to dictate the desired pillar spacings of 5, 10, 20, and 50 µm. To create the positive template, a silicon wafer was cleaned with 100% ethanol and dried. A thin layer of SU-8 photoresist (Fisher Scientific; NC1248659) was then spin coated onto the wafer. Using a Karl Suss MA-4 Mask Aligner, the mask was placed over the wafer and both were exposed to UV light with an exposure of 25 seconds to crosslink the photoresist at the designated pillar locations. Excess photoresist was dissolved using SU-8 developer. The finished positive template was silanized overnight in a vacuum desiccator with tridecafluoro-1,1,2,2,tetrahydrooctyl-1-trichlorosilane (OTS, 97%) (TCI Chemicals; T2577-5G). Pillar height was confirmed using a profilometer to ensure that all pillars were within a 14-17 µm range. All positive template fabrication was done in the University of Maryland Nanocenter FabLab.

The silanized positive template was then used to create a negative template made of PDMS. Sylgard 184 PDMS was mixed at a 10:1 base to crosslinker weight ratio and 50 g was poured onto the positive template. Air bubbles were removed by placing the dish into a vacuum desiccator for 20-30 minutes, and the PDMS was cured at 80°C for 1 hour. The cooled PDMS was carefully peeled off the wafer and the negative template was cleaned off with filtered air, plasma treated for 3 minutes, and silanized overnight in a vacuum desiccator with tridecafluoro-1,1,2,2,tetrahydrooctyl-1-trichlorosilane (OTS, 97%).

To fabricate the final micropillar devices, 12 g of the mixed PDMS was poured onto the negative template, degassed in the vacuum desiccator for 20-30 minutes, and cured at 80°C for 1 hour. Cooled PDMS was carefully peeled off and the devices were cut out. Each micropillar device was kept whole for use with the bioreactor or cut into quarters for the static culture groups.

One day prior to device functionalization, pluronic stamps were prepared by pouring 8% Pluronic F-127 (Sigma-Aldrich; P2443-250G)) onto dishes of Sylgard 184 PDMS which were then Parafilmed and left to incubate at room temperature overnight.

The next day (one day prior to seeding), devices were washed with alternating 100% ethanol and DI water and dried with filtered air and placed in either 6-well plates or 100 x 15 mm petri dishes. For the stamps, pluronic was poured off and the surface was rinsed with 100% ethanol and DI water before being dried with filtered air. Devices were plasma treated for 6-7 minutes, gently placed pillar side-down on the prepared stamps for 5 minutes, and then UV sterilized for 10 minutes. The devices were submerged in a 20 µg mL^-1^ collagen I solution for at least 1 hour at 37°C. The collagen was then removed, devices were washed once with 1X PBS and seeded with either 25,000 cells for the quartered devices, or 100,000 cells for the whole devices. For the flask + collagen groups, T-175 flasks were coated with 20 µg mL^-1^ of collagen I and seeded with 250,000 cells. The next day, media was replaced with EV-depleted iMSC media for the static confinement and flask groups. For the bioreactor group, further setup as detailed below was followed.

### 3D-printed bioreactor fabrication and setup

Bioreactor design parameters were altered from a previous design from our group [8] to have a flat interior and removable lid to allow for implantation of the micropillar devices. SolidWorks designs were exported into Envison One RP software for printing. Bioreactors were 3D-printed with LOCTITE Med413 resin from ETEC, using an ETEC Envision One DLP printer. Bioreactors were submerged in an isopropyl alcohol bath following manufacturer’s recommendations to wash away excess resin. Complete crosslinking was achieved by two rounds of 20-minute UV exposure.

Prior to use, 3D-printed bioreactors were rinsed with 100% ethanol and dried with filtered air before UV sterilizing for 10 minutes. Seeded PDMS micropillar devices were inserted into the center of the bioreactor base using super glue. The lid was fitted with a rubber O-ring and placed onto the base, which was then connected to a Masterflex L/S Digital Drive (Cole-Parmer) coupled with a pump head to perfuse media at flow rates of 1, 5, or 10 mL min^-1^. Both open ends of the tubing were placed in a reservoir of 70 mL of EV-depleted iMSC media, while the other ends connected to the inlet and outlet of the bioreactor. The assembled bioreactor-pump system was placed in an incubator at 37°C with 5% CO_2_ and allowed to run for 24 hours. Bioreactors were not reused.

### EV separation

Conditioned media was collected from the static confinement devices and flasks and replaced with fresh EV-depleted media for 3 days. Bioreactors were run for 24 hours, and the conditioned media was collected. All media was subjected to three differential centrifugation spins (1000 x *g* for 10 minutes, 2000 x *g* for 20 minutes, and 10,000 x *g* for 30 minutes) and then passed through a 0.2 µm filter. Final EVs were isolated using tangential flow filtration (TFF) (Repligen; KrosFlo KR2i TFF system) using a protocol adapted from Heinemann et al. [58]. Briefly, samples were concentrated down to 10-15 mL using a 100 kDa membrane filter (Repligen; D04-E100-05-N) and subjected to 10 diafiltration steps and a transmembrane pressure of 5 psi. Samples were then concentrated further using a 100 kDa centrifugal concentrator (Corning; 431486) and resuspended in 1X PBS.

### EV characterization

EV concentration and size was determined using nanoparticle tracking analysis (). Three 30 second videos were acquired for each sample, and camera levels and detection threshold were maintained between all batches. Total protein concentration was quantified using a bicinchoninic acid (BCA) assay per the manufacturer’s protocol (G-Biosciences; 785-571).

EV identity was confirmed using Western blotting with equal amounts of protein or cell lysate per lane. Samples were run through the gels and transferred to nitrocellulose membranes. EV-positive markers CD63 (ThermoFisher Scientific; 25682-1-AP), ALIX (Abcam; ab186429), TSG101 (Abcam; ab125011), and EV-negative marker calnexin (Cell Signaling Technology; 2679) were probed. All primary antibodies were incubated overnight at 4C and diluted at a 1:1000 dilution. The next day, goat anti-rabbit IRDye 800CW (LI-COR; 925-32210) was incubated with the PBS-washed membranes for 1 hour at a 1:10,000 dilution. Membranes were imaged using an Odyssey CLx imager (LI-COR).

Transmission electron microscopy (TEM) images of the EVs were captured using a negative stain. EV samples were fixed in 4% paraformaldehyde (PFA) for 30 minutes at room temperature. A carbon film grid (Electron Microscopy Sciences; CF-200-Cu-25) was placed on a droplet of the EV/PFA solution for 20 minutes, then washed with a droplet of 1X PBS followed by 1% glutaraldehyde in PBS for 5 minutes. The grid was again washed using a droplet of DI water 5 times for 2 minutes each and then placed on a droplet of uranyl acetate replacement stain for 10 minutes (Electron Microscopy Sciences; 22405). Grids were allowed to dry and imaged using a JEM 2100 LaB6 TEM (JEOL USA Incorporated).

### EV production quantification

For EV production data, conditioned media was collected only once. After collection, the cells in the confinement devices were fixed using 4% PFA for 30 minutes on ice on a shaker. PFA was aspirated and cells were then washed three times with 1X PBS before adding a 0.5% Triton X-100 in PBS solution for 7 minutes to permeabilize the membranes. Triton X-100 was aspirated and cells were again washed three times with 1X PBS. 2 µg mL^-1^ DAPI was added and incubated for 30 minutes in the dark and then aspirated. Cells were washed once with 1X PBS and devices were turned upside down and imaged using an IX-83 microscope. Total cells were quantified using ImageJ. For flask cultures, cells were trypsinized and counted. The total number of EVs per sample was determined using NTA as described above, and EV production as measured by EV per cell was calculated.

### Tube formation assay

In a 48-well plate, wells were coated with 100 µl of growth factor reduced Matrigel (Corning; 356230) and incubated for at least 30 minutes at 37°C. 35,000 P3 or P4 HUVECs or HDMECs were mixed with either endothelial growth media with 5% FBS (positive control), endothelial basal media with 0.1% FBS (negative control), or 5×10^9^ EVs mL^-1^ in basal media. 4-6 hours later, tubes were imaged using a Nikon Eclipse Ti2 Microscope at 2x magnification and number of fully formed tubes were counted.

### Gap closure assay

96 well plates were coated with 0.1% gelatin for at least 1 hour at 37°C for assay with HUVECs. No coating was necessary for fibroblasts and keratinocytes. 60,000 cells per well of P3 or P4 HUVECs, or 40,000 cells per well of P3 or P4 fibroblasts or keratinocytes were seeded and left to adhere overnight to form a monolayer. The next day, the respective growth medias were replaced with their basal media counterparts ∼2 hours before applying EV treatments. A scratch was then induced using a p200 pipette tip, the wells were washed with 1X PBS, and either growth media, basal media, or 5×10^9^ EVs mL^-1^ in basal media were added. Scratches were imaged immediately after and again 20 hours later at the same location. Percent gap closure was quantified using ImageJ. HUVEC growth and basal media consisted of endothelial growth media with 5% FBS and 1% penicillin-streptomycin and endothelial basal media with 0.1% FBS and 1% penicillin-streptomycin, respectively. Fibroblast growth and basal media consisted of DMEM/F-12 with 10% FBS and 1% penicillin-streptomycin and DMEM/F-12 with 0.1% FBS and 1% penicillin-streptomycin, respectively. Keratinocyte growth and basal media consisted of dermal cell basal media supplemented with ATCCs’ keratinocyte growth kit components and 0.1% penicillin-streptomycin and dermal cell basal media with 0.1% penicillin-streptomycin, respectively.

### Anti-inflammatory assay

RAW 264.7 mouse macrophages were seeded at 75,000 cells per well in a 48-well plate in DMEM with 5% FBS and 1% penicillin-streptomycin. 24 hours later, cells were pre-treated with either 5×10^9^ EVs mL^-1^ or left untouched (positive control). The following day, media was replaced with 10 ng mL^-1^ lipopolysaccharide (LPS) in media and left to incubate for 4 hours at 37°C. The conditioned media was then collected and stored at -80°C. Pro-inflammatory cytokines IL-6 and TNF-⍰ were measured using DuoSet ELISA kits (R&D Systems; DY406, DY410).

### Molecular content analysis

For the miRNA qPCR arrays, EVs were lysed using QIAzol Lysis Reagent (Qiagen; 79306) and RNA was isolated using the Zymo Direct-zol RNA Microprep kit (Zymo Research; R2061) and reverse transcribed using the All-in-One miRNA First-Strand cDNA Synthesis Kit (GeneCopoeia; QP018). Resulting cDNA was analyzed in a human MSC exosome miRNA qPCR array (GeneCopoeia; QM0460B6) per the manufacturer’s protocol. Quantitative polymerase chain reaction (qPCR) was performed using a QuantStudio 7 Flex qPCR system (ThermoFisher Scientific; 4485701), and data were analyzed using the delta-delta Ct method normalized to the housekeeping gene SNORD48.

For proteomics, proteins were isolated from EV samples using the Universal Proteomics Sample Prep Kit from ProtiFi (ProtiFi; K02-Micro-10) using the provided buffers and following the manufacturer’s protocol. Briefly, the sample was lysed and solubilized. The disulfide bonds were then reduced and alkylated, and the proteins were denatured. The sample, with the specified buffer, was added to the S-Trap column and centrifuged to allow the proteins to be separated from the solution. The sample was then washed 5 times to remove contaminants. 2 µg trypsin was added to the column and incubated overnight at 37°C in a humidified chamber to digest the proteins. The next day, peptides were eluted, dried down, and resuspended in 5% acetonitrile and 0.1% formic acid. Proteomics analysis was run by the Molecular Characterization and Analysis Complex at the University of Maryland Baltimore County using the timsTOF Pro 2 (Bruker) coupled with the nanoElute HPLC (Bruker) for sample separation. Results were analyzed with PEAKS Studio software by Bioinformatics Solutions.

Gene ontology/biological process pathway enrichment analysis was performed using the PANTHER 19.0 online database system [31–33]. Fisher’s exact test with false discovery rate (FDR) correction was used for analysis.

### Diabetic mouse wound healing model

All animal care, experiments, and handling were performed in accordance with NIH, local, state, and federal guidelines and approved by the Institutional Animal Care and Use Committee (IACUC) (protocol R-AUG-23-41) at the University of Maryland College Park.

18 male Lepr^db^ (db/db) mice at 8-12 weeks of age from Jackson Laboratory were used in this study. Mice were initially anesthetized using 3% isoflurane, which was then switched to 1.5% isoflurane once they were moved to the nose cone to maintain sedation. 2.5 mg kg^-1^ Banamine was administered subcutaneously, the entire dorsum was shaved, and the area was cleaned three times with alternating swabs of 70% ethanol and iodine. An 8 mm punch biopsy (Medical Device Depot; 33-37) was then used to create a full thickness wound in the center of the dorsum. 1.1×10^10^ total EVs in 200 µL PBS were injected intradermally in a cross pattern around the wound with a total of 4 injections (2.75×10^9^ EVs per 50 µL injection). An equivalent volume of PBS was used for the control group. A silicone ring with an inner diameter of 12 mm and an outer diameter of 17 mm was adhered to the skin around the wound using Dermabond (Medical Supply 123; DNX12) and simple interrupted sutures. Images were taken and the wound was covered with Tegaderm. One day later, 2.5 mg kg^-1^ Banamine was again administered subcutaneously. EV or PBS injections were repeated on days 3, 6, and 9, and images were also taken to assess wound closure. On day 16, mice were euthanized, and healed wound tissue was biopsied using a 12 mm punch biopsy (Acuderm; P1225).

After biopsy, the tissue was cut down the middle and half was placed in RNAlater (ThermoFisher Scientific; AM7020) for future RNA isolation. The other half was fixed in 10% neutral buffered formalin for 24 hours at 4°C before being transferred to 70% ethanol for embedding, 5 µm sectioning, mounting, and H&E staining which was performed by the University of Maryland School of Medicine’s Pathology Histology Core. Masson’s Trichrome staining. All slide digitization and H&E and Masson’s Trichrome interpretation was performed by the University of Maryland School of Medicine’s & Greenebaum Comprehensive Cancer Center’s Pathology Biorepository Shared Resources Core.

For CD31 IHC, mounted slides were rehydrated and antigen retrieval was performed by heating slides in a 10mM sodium citrate solution (pH 6.0) at 95°C for 10-15 minutes. Slides were cooled in DI water for 2 minutes and tissue sections were circled with a hydrophobic barrier PAP pen (Vector Laboratories; H-400). Sections were washed with PBS and then blocked with a solution of PBS with 5% donkey serum (Sigma-Aldrich; D9663-10ML), and 1% BSA (GoldBio; A-421-100) for 30 minutes in a humidified chamber. Excess liquid was removed and slides were incubated with CD31 antibody (Abcam; 28364) diluted 1:50 in blocking solution for 1 hour. Slides were washed twice with PBS for 5 minutes and incubated with 5 µg mL^-1^ Alex Fluor 647 donkey anti-rabbit secondary antibody (ThermoFisher Scientific; A31573) in a humidified chamber for 1 hour. Slides were again washed twice with PBS, and coverslips were mounted after adding Vectashield Mounting Medium (Vector Laboratories; H-1200-10). Coverslips were sealed with clear nail polish and fluorescence images were taken on a FV3000 Laser Scanning Confocal Microscope (Olympus) at 10X magnification over multiple fields of view for each tissue section, keeping all laser settings consistent between samples. Fluorescence intensity for each channel was quantified with ImageJ where the threshold settings were maintained between all images.

Tissue stored in RNAlater was homogenized in QIAzol using a Scilogex D160 Homogenizer and centrifuged at 20,000 x *g*. Supernatant was extracted and RNA was extracted using the Zymo Direct-zol RNA Miniprep kit (Zymo Research; R2050) and then reverse transcribed using the RevertAid First Strand cDNA Synthesis Kit (ThermoFisher Scientific; K1622). qPCR for M1 and M2 markers was performed using GAPDH as a housekeeping gene and run on a QuantStudio 7 Flex qPCR system (ThermoFisher Scientific; 4485701). Data were analyzed using the delta-delta Ct method.

### Statistical analysis

All data is represented as the mean +/- standard error of the mean. An ordinary one-way ANOVA with Sidak’s multiple comparisons test, two-way ANOVA with Holm-Sidak’s multiple comparison test, or unpaired t-tests with Holm-Sidak’s multiple comparisons test was used to determine statistical significance. Statistical analyses were performed using Prism 10 (GraphPad Prism Software). Statistical significance is indicated as: ns (p > 0.05), * p<0.05, ** p<0.01, *** p<0.001, **** p<0.0001.

## Acknowledgements

The authors would like to acknowledge the University of Maryland School of Medicine’s Pathology Histology Core and the University of Maryland School of Medicine’s and Greenebaum Comprehensive Cancer Center’s Pathology Biorepository Shared Resources-Baltimore, MD. The Pathology Biorepository Shared Resources was supported by funds through the Maryland Department of Health’s Cigarette Restitution Fund Program and the NCI-Cancer Center Support Grant P30CA134274. The authors would also like to acknowledge the Molecular Characterization and Analysis Complex at the University of Maryland Baltimore County.

## Data availability statement

The data that supports the findings of this study are available from the corresponding author upon reasonable request.

