## Supporting Information for "Perfusion Bioreactor Culture Incorporating Mechanical Confinement Enhances Mesenchymal Stem Cell Extracellular Vesicle Production and Wound Healing Potential"

### SUPPLEMENTAL DATA

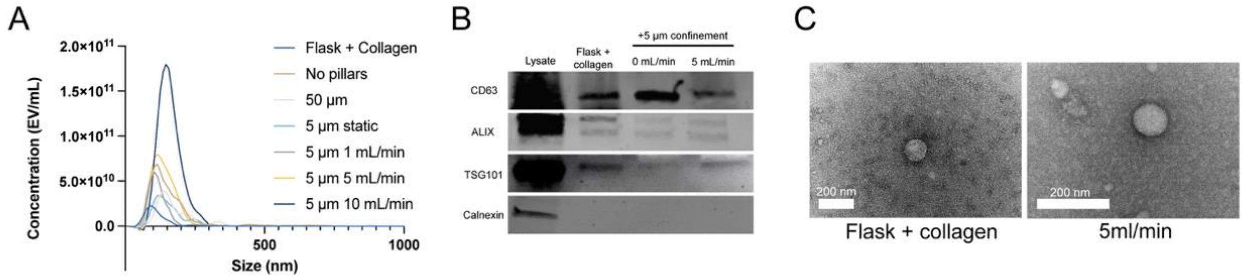


**Figure S1: iMSC EV characterization.** A) EV size and concentration profiles generated via nanoparticle tracking analysis. B) Representative western blots of EV samples, probing for EV markers CD63 (25-65 kDa), ALIX (95 kDa), and TSG101 (44 kDa) and EV-negative marker calnexin (90 kDa) with an equal amount of total protein loaded per lane. C) Representative transmission electron microscopy images of EVs from cells in flask culture and confined micropillar culture.

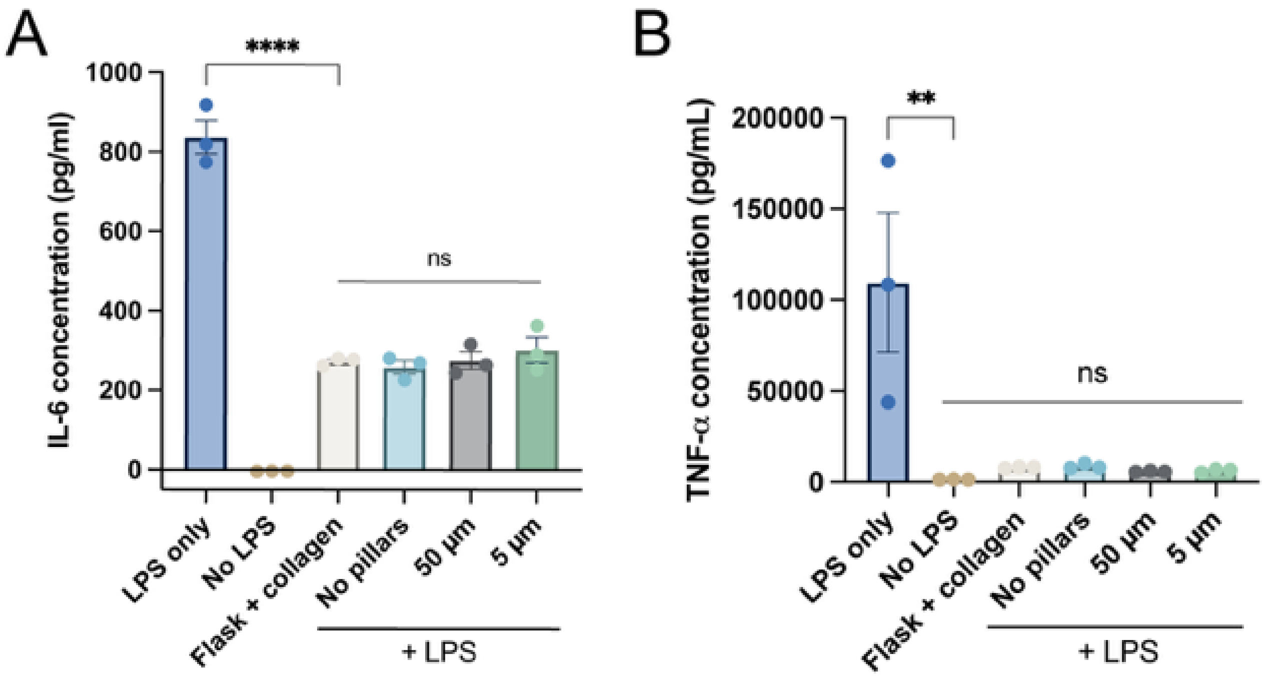


**Figure S2: Anti-inflammatory effects of EVs from confined producer cells.** A) IL-6 and B) TNF-α production by RAW264.7 mouse macrophages after stimulation with LPS and treatment with 5E9 iMSC EVs/mL from different static culture conditions with increasing confinement, as measured by ELISA (n=3). All data is representative of 3 independent experiments. All values expressed as mean +/- standard error of the mean (ns – no significance; ** $p<0.01$; **** $p<0.0001$).

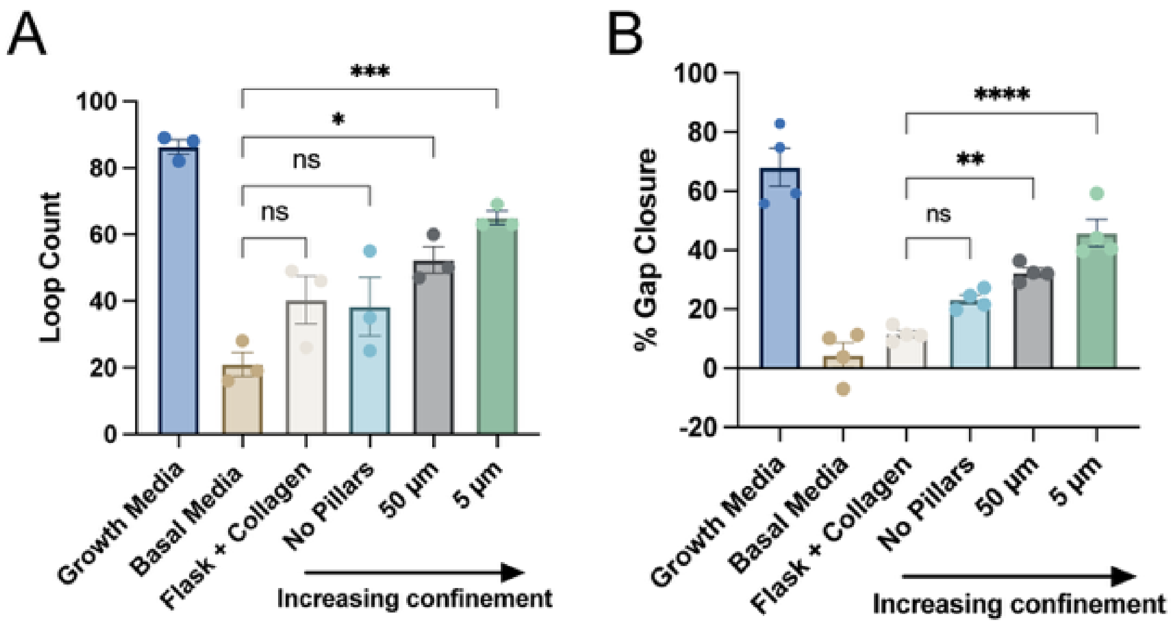


**Figure S3: BMMSC EV bioactivity is responsive to producer cell confinement.** A) Tube formation and B) percent gap closure data by HUVECs treated with 5E9 bone marrow-derived MSC EVs/mL from different culture conditions with increasing levels of confinement (n=3). All data is representative of 3 independent experiments. All values expressed as mean +/- standard error of the mean (ns – no significance; * $p<0.05$; ** $p<0.01$; *** $p<0.001$; **** $p<0.0001$)

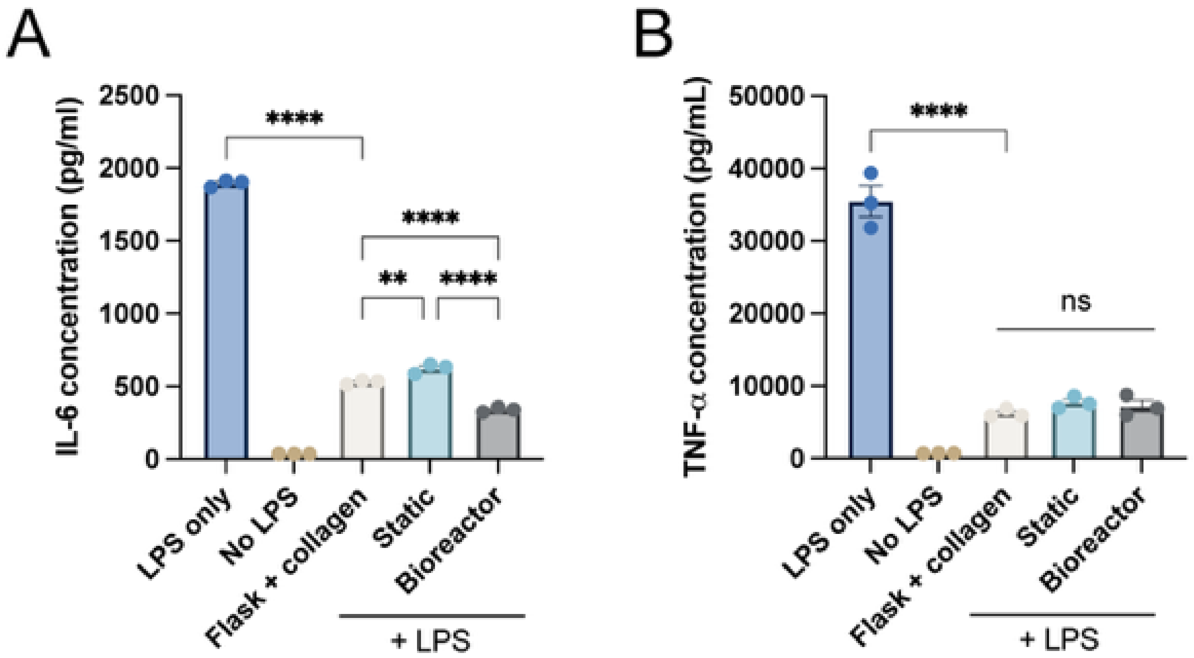


**Figure S4: Anti-inflammatory effect of EVs from confined cells in static and optimized bioreactor culture.** A) IL-6 and B) TNF-α production by RAW264.7 mouse macrophages after stimulation with LPS and treatment with 5E9 iMSC EVs/mL from flask, static confinement, and bioreactor confinement culture, as measured by ELISA (n=3). All data is representative of 3 independent experiments. All values expressed as mean +/- standard error of the mean (ns – no significance; ** $p<0.01$;**** $p<0.0001$).
